# Microbial dispersal limitation to isolated soil habitats in the McMurdo Dry Valleys of Antarctica

**DOI:** 10.1101/493411

**Authors:** Stephen D.J. Archer, Kevin C. Lee, Tancredi Caruso, Teruya Maki, Charles K. Lee, Don A. Cowan, Fernando T. Maestre, Stephen B. Pointing

## Abstract

Dispersal is a critical yet poorly understood factor underlying macroecological patterns in microbial communities. Airborne microbial transport is assumed to occupy a central role in determining dispersal outcomes and extra-range dispersal has important implications for predicting ecosystem resilience and response to environmental change. One of the most pertinent biomes in this regard is Antarctica given its geographic isolation and vulnerability to climate change and human disturbance. Here we report the first characterisation of microbial diversity in near-ground and high-altitude air above a typical Antarctic Dry Valley as well as that of underlying soil microbial communities. We found that persistent airborne inputs were unable to fully explain local soil community assembly. Comparison with airborne microbial diversity from non-polar sources suggests that strong selection occurs during atmospheric transport resulting in regionally isolated airborne inputs and highly specialized soil communities where fungi displayed greater isolation than bacteria from non-polar sources. Overall microbial communities from this isolated Antarctic ecosystem displayed limited connectivity to the global microbial pool. Our findings provide critical insights to forecast the potential outcomes for microbial communities of climate change-mediated shifts in air circulation to the Dry Valleys, the largest ice-free region of Antarctica.

## Introduction

A key determinant of biogeographic and macroecological patterns in all microbial communities is dispersal ^1^. Microbial uptake to the aerosphere, persistence during transit and deposition all play a central role in determining dispersal outcomes ^2,3^ and extra-range dispersal has important implications for predicting ecosystem resilience and response to environmental change ^4^. Airborne microbial transport on fine particulate matter has typically been regarded as ubiquitous due to the small size and survivability of cells ^5–7^. Therefore, large scale patterns in microbial diversity are often viewed as developing largely due to deterministic niche-driven processes ^8,9^. Much of the existing evidence for airborne (also known as aeolian) transport of microorganisms, typically viewed as a neutral process ^10^, is inferred from extant communities at source and sink locations e.g. ^11–13^. Furthermore, direct measurements of airborne microbial taxa have largely focused on indoor or outdoor built environments e.g. ^14–16^ rather than across natural environmental gradients. This limits our understanding of how microbial transport drives biogeographic patterns. The extent to which the aerosphere is a habitat as opposed to simply a medium for microbial transport is also unresolved and as a result the influence of airborne transport on observed patterns in microbial biogeography has been subject to much speculation supported with little tangible evidence ^17–19^.

Antarctica is a focus for microbial ecology research due to its geographic isolation, lack of trophic complexity and human influence ^20^ and vulnerability of endemic biodiversity to climate change ^21^. The extent to which airborne immigration may influence isolated Antarctic terrestrial microbial communities, however, is an enduring enigma in microbial ecology ^22–25^. Highly specialised microbial communities display strong allopatric signals ^26–29^, yet microbial dispersal is conventionally viewed as occurring across inter-continental distances ^22,30^. Establishing the extent to which Antarctic communities are connected to the global system via airborne dispersal also has key relevance to predicting potential responses of polar ecosystems to environmental change. Isolated ice-free regions such as the McMurdo Dry Valleys are devoid of vascular plants and dominated by highly specialised soil microbial communities ^31^ that display adaptations to the extreme environmental conditions ^32^. Some taxa, notably cyanobacteria, have been shown to display phylogenetic endemism in Antarctica at the level of rRNA gene-defined diversity ^26–29^. Also, lichenised fungi displayed patterns in diversity that suggest that they radiated from local refugia rather than from exogenous sources outside Antarctica ^33^. A global theoretical model for atmospheric aerosols estimated that the rate of airborne microbial exchange to Antarctica may be extremely low, with 90% of aerosols expected to be of local origin ^34^. In contrast, empirical studies have claimed circum-polar distribution for some Cyanobacteria, chlorophyte algae, and Fungi ^22,30,35^. Antarctica therefore presents a paradox in microbial biogeography with regard to microbial dispersal.

Evidence for airborne microorganisms in Antarctica is scarce: two early studies on the Antarctic Peninsula (the west continental edge of Antarctica proximal to South America) identified tentative evidence for airborne bacteria from individual samples without characterisation beyond a low diversity of common taxa and human- and penguin-associated bacteria ^36,37^. We recently observed that in the relatively isolated ice-free McMurdo Dry Valleys soil region of East Antarctica airborne bacteria were recoverable from bulk air and may harbour far greater diversity than previously envisaged ^38^. These studies were generally inconclusive as to the origin and relationship to local habitats due to methodological constraints (Online Supplementary Material). Here we use state-of-the art sampling, sequencing and statistical approaches to study the diversity of airborne and soil microbial communities in a typical Antarctic Dry Valley. We targeted Bacteria and Fungi because these domains are the most abundant microorganisms in the McMurdo Dry Valleys ^31^. We tested the null hypotheses that air and soil microbial communities are a random sample of the regional and global microbial pools. For doing so, we acquired massive bulk-air samples and estimated, for the first time, microbial diversity in near-ground air and underlying soil for low and high elevation sites, as well as polar air above the boundary layer for surface interactions and non-polar sources.

## Results and Discussion

The incoming air mass to our study site in the McMurdo Dry Valleys largely transited above the Antarctic Plateau during the maximum predicted residence time for bacteria and fungi in air (15 days) ^34^, whilst the most distant air mass had a non-polar origin above the coastal shelf of New Zealand (Fig. 1a). Transport was exclusively from the Polar Plateau and across the Trans-Antarctic Mountains during the average residence time for microorganisms in air (Fig. 1a). We thus envisage that severe selection pressure should occur during airborne transit in an air mass with freezing temperatures and high UV exposure at mean altitudes of 2769m (3 day transit) and 3034m (15 day transit).

**Fig. 1.**
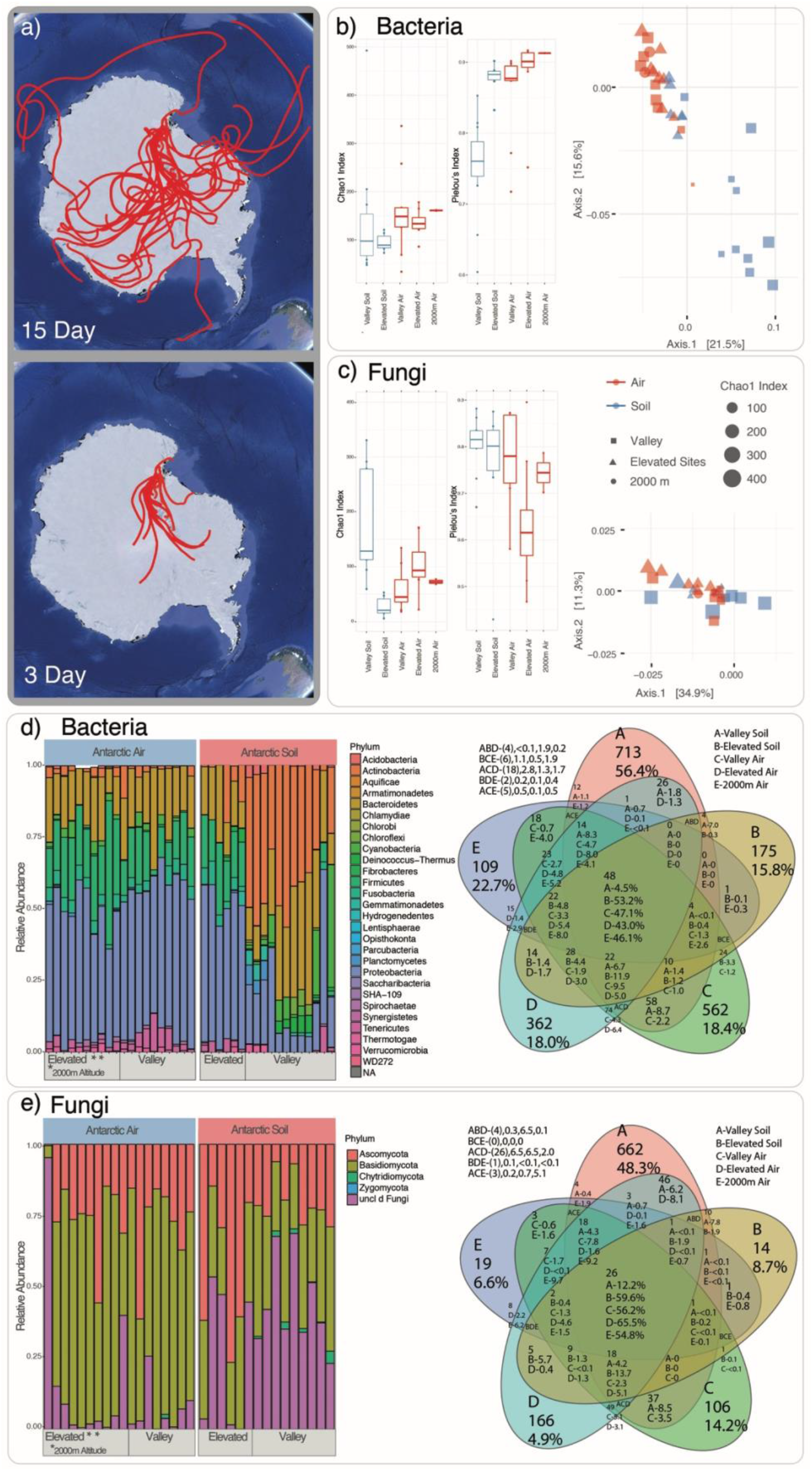
Antarctic air and soil habitats support distinct bacterial and fungal communities. **a)** Route predictions for average minimum and maximum modelled residence time for microorganisms in air based on HYSPLIT back trajectory analyses. Back trajectories indicate distance travelled for sampled air mass at 3 d (598 - 2581 km distance, average altitude of 2769 m, maximum altitude of 5174 m A.M.S.L.) and 15 d (4673 - 11216 km distance average altitude 3034m, maximum altitude 6886 km). **b) and c)** Alpha diversity estimates (Chao1 richness and Pielou’s relative evenness) and visualisation of community dissimilarity using Principal Co-ordinate Analysis of weighted UniFrac distance by habitat for **b)** Bacteria and **c)** Fungi. Boxplot whiskers represent 1.5 times the interquartile range from the first to the third quartiles or the maximum/minimum data point within the range. **d) and e)** Distribution and relative abundance of **d)** Bacteria and **e)** Fungi in Antarctic air and soil. Each stack bar represents data from three pooled replicates for each substrate location. Diversity is shown at phylum level as this is the highest taxonomic rank at which between-substrate differences are noticeable. Venn diagrams show amplicon sequence variants (ASVs) and operational taxonomic units (OTUs) count and percentage occurrence within and between each habitat. The high altitude samples, i.e., those without underlying soil, do not have corresponding soil samples. Sampling locations: valley (soil and 1.5m above ground), elevated (soil and 1.5m above ground at higher altitude locations at Bull Pass and valley ridges), 2000m (helicopter samples). Interactive graphics identifying taxonomic composition to lower taxonomic ranks within each sample are presented in the Online Supplementary Information (Fig. S2). Comparison with non-polar samples is given in the Online Supplementary Information Fig. S2.

In general terms, we found that alpha diversity metrics for air and elevated high-altitude soils were distinct from those observed in valley soils, with taxa richness being more variable in valley soils than in air and elevated soils (Fig. 1b,c). These results highlight the more heterogenous nature of valley soils as a habitat compared with bulk phase air or elevated altitude soils, where conditions are generally unfavourable to colonisation. The elevated altitude mineral soil sites may therefore be representative of near-term airborne deposition to this system. Bacterial taxa richness was similar among all samples although slightly higher in air and elevated soil (Fig. 1b). Conversely, the richness of fungal taxa was highest in valley soils, and lower in air and elevated soils (Fig. 1c). Soils also displayed greater evenness and lower richness values than air samples and this tentatively indicated that airborne fungi are under strong selective pressure. Ordination analyses of weighted UniFrac distances for bacteria and fungi supported these trends in alpha diversity (Fig. 1b,c). Valley soil bacterial communities separated clearly from air and elevated soil communities (Fig. 1b). A similar although less pronounced pattern was observed for fungi (Fig. 1c).

Taxonomic assignment of bacteria and fungi revealed further complexity. Airborne phylum-level bacterial diversity was dominated by Proteobacteria, Bacteroidetes and Firmicutes (Fig. 1d, Online Supplementary Information Fig. S2). Phyla with high relative abundance comprised spore-formers and taxa with known UV and/or desiccation tolerant traits viewed as advantageous during atmospheric transport ^17^ and survival in Antarctic soil ^32^. The airborne bacterial samples supported relatively high levels of taxa associated with marine influence ^7^ suggesting recruitment during transit over the Southern Ocean (Online Supplementary Information, Fig. S2). Terrestrial bacteria were also transported and may have benefited from islands acting as stepping stones for dispersal ^7^. Soil communities supported greater abundance of Actinobacteria and other taxa typical from arid soils ^8^. Near-ground air supported 3-5 fold more habitat-specific taxa than high-altitude air; samples from the former habitat were most similar to their underlying soil communities. Valley soils supported 56.4% soil-specific taxa compared with only 15.8% in elevated high-altitude soils. Valley soils shared very few taxa with the total air sample pool (4.5%) whilst different air habitats (valley, elevated and high-altitude) shared approximately half the taxa encountered in each habitat.

The most abundant fungal taxa in air were basidiomycetous yeasts (Fig. 1e, Online Supplementary Information Fig. S2) whereas soils were dominated by unclassified fungi also including yeasts. The yeasts are thought to be well-adapted to growth in Antarctic soil habitats ^39^. Ascomycetes were also commonly encountered but Chytrids occurred only in valley soil and air. Valley soils supported 48.3% habitat-specific taxa while elevated soils and high-altitude air showed the lowest number of habitat-specific taxa (4.9-8.7%). The different air habitats shared approximately half the taxa encountered in each habitat. Fungi are well-adapted to conditions anticipated during atmospheric transport due to the production of resistant spores and UV-protective compounds ^40,41^. Local recruitment may, however, be limited to asexual states since teleomorph fruiting structures are not known from Antarctic fungi ^42^.

We achieved near-asymptote in diversity estimation for all samples (Online Supplementary Information, Supplementary Methods). Thus we further interrogated the phylogenetic diversity of air and soil by generating distribution heatmaps by habitat for the 1,000 most abundant taxa. This analysis captured 91% of total bacterial and 96% of total fungal diversity in the libraries (Fig. 2a,b). We also incorporated diversity data from air originating at the nearest non-polar land mass into this analysis. A striking pattern emerged where soil bacterial and fungal assemblages in the McMurdo Dry Valleys were only partially recruited from local air taxa. This pattern cannot be further explained by airborne recruitment from exogenously sourced aerosols. These findings support the notion of a system operating in stark contrast with the long-held assumption that microbial dispersal is ubiquitous and deterministic niche processes are the primary driver of community assembly in terrestrial surfaces ^1,10^. We therefore further interrogated this association using Ecological Network Analysis (Fig. 2c). Overall non-polar air displayed least connectivity to all other Antarctic habitats as previously observed by the weak associations and greater Bray-Curtis distances between them (Fig. 2c). Bacterial communities clustered by habitat type (Fig. 2c). This pattern is likely to be indicative of selection pressures due to local environmental filtering, which could combine a mixture of biotic and abiotic factors. Conversely, fungal communities associated by geographic distance and were thus more likely to be influenced by dispersal limitation. No significant distance-decay relationships were observed for airborne or soil communities between valley and elevated sites (air bacteria R^2^ = 0.009, soil bacteria R^2^ = 0.006, air fungi R^2^ = 0.016, soil fungi R^2^ = 0.024). These results indicate that dispersal within the Dry Valleys may be limited and likely reflects the associated steep environmental gradients present in this region. These findings provide empirical support to prevailing theoretical models of emission and transport for biological particles in the atmosphere that predicts relatively low exchange between Antarctic and non-polar air as well as reduced residence time in air for fungi compared to bacteria due to allometric considerations ^34^.

**Fig. 2.**
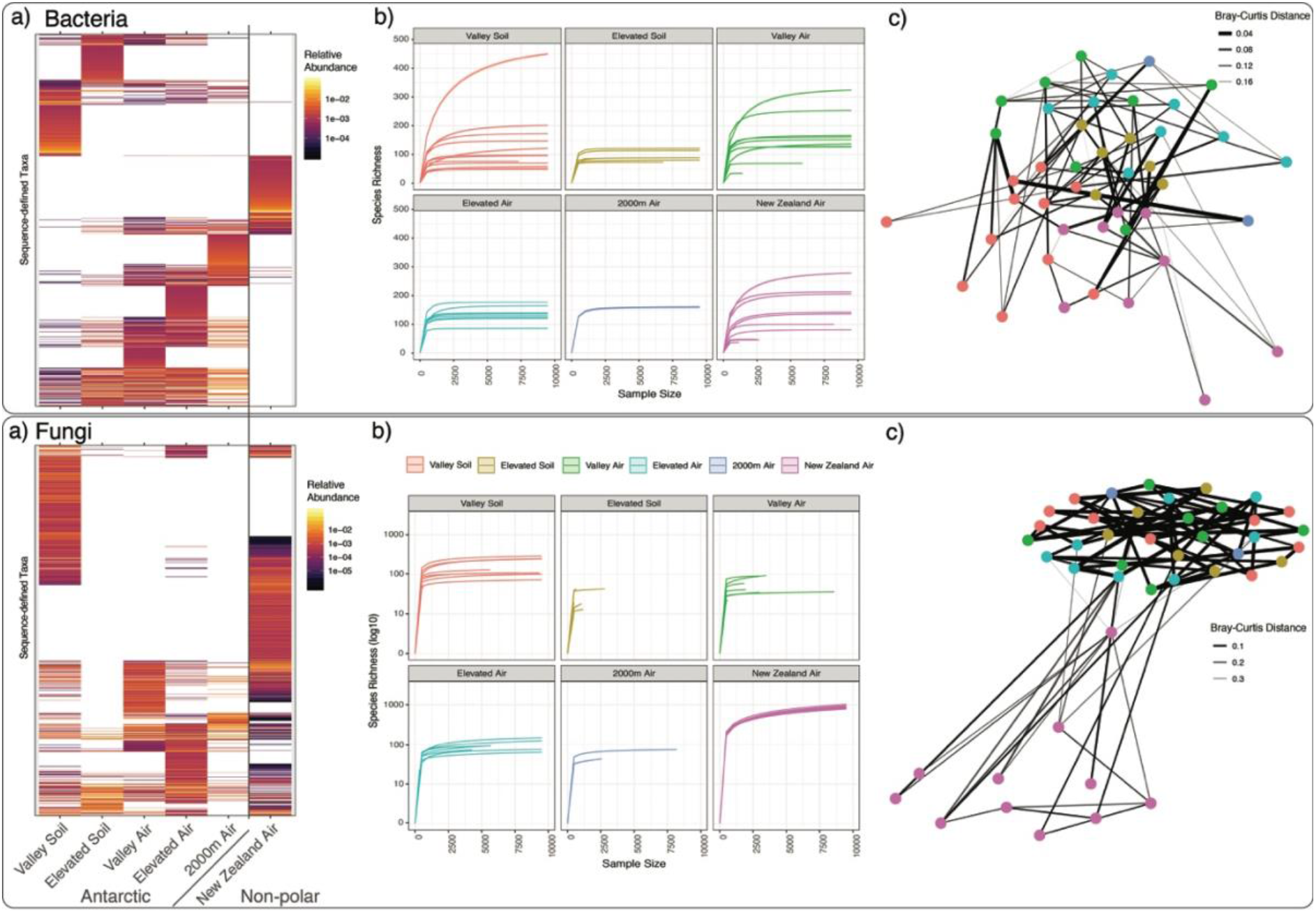
Comparison of bacterial and fungal diversity from Antarctic and non-polar sources. **a)** Distribution and relative abundance for the 1,000 most abundant bacterial amplicon sequence variants (ASVs) and fungal operational taxonomic units (OTUs). **b)** Rarefaction curves are shown for Bacteria and Fungi for each Antarctic habitat to illustrate sampling depth to near-asymptote. **c)** Co-occurrence associations derived from Ecological Network Analysis. We enforced maximum Bray-Curtis distance of 0.2 for Bacteria and 0.4 for Fungi to establish connection between nodes (representing communities). The nodes were positioned using the Fruchterman-Reingold method. Sampling locations: valley (soil and 1.5m above ground), elevated (soil and 1.5m above ground at higher altitude locations at Bull Pass and valley ridges), 2000m (helicopter samples) and New Zealand (non-polar). Full taxonomic comparison for all polar and non-polar samples is given in the Online Supplementary Information Fig. S2.

We conducted additional analyses (Nestedness Analysis and Net Relatedness Index analysis) to reveal the extent to which taxonomic and phylogenetic structuring reflected the likelihood of exogenous recruitment (Fig. 3). Strong patterns of nestedness are the classical expectation for a network of highly isolated sites (e.g., distant islands) where passive sampling from regional pools and ordered sequences of extinctions play the major roles in structuring local communities ^43,44^. Strong or perfect nestedness indicates that species poor local communities are a proper subset of richer communities. We used one of the most widely applied metrics of nestedness (NODF ^44–46^) and applied it to the occurrence of the 1,000 most abundant bacterial and fungal taxa. The NODF metric, which ranges from 0 (no nestedness) to 100 (perfect nestedness), can be decomposed into a compositional effect which ranges from 0 (no nestedness) to 100 (perfect nestedness). This illustrates a compositional effect (i.e., species poor communities consist of species that are a subset of richer communities; NODFc) and an incidence effect (i.e., less frequent species always occur in site with widespread species; NODFr). Null models applied to these metrics showed that communities of both Bacteria and Fungi were significantly anti-nested (NODF <30 approx.) (Fig. 3a). This general result implies that passive sampling from the regional pool alone is not sufficient to explain the structure of local Antarctic communities. Both Bacteria and Fungi, however, were significantly nested for taxonomic composition under the hypothesis that nestedness can be maximised by ordering sites from the most connected, to the least connected, to a global species pool. This suggests that species poor assemblages of the least connected sites are a proper subset of richer, more connected sites. Fungi were markedly more nested (NODFc = 62) than Bacteria (NODFc = 18) (Fig. 3a), suggesting a potential major role of dispersal limitation for this group.

**Fig. 3.**
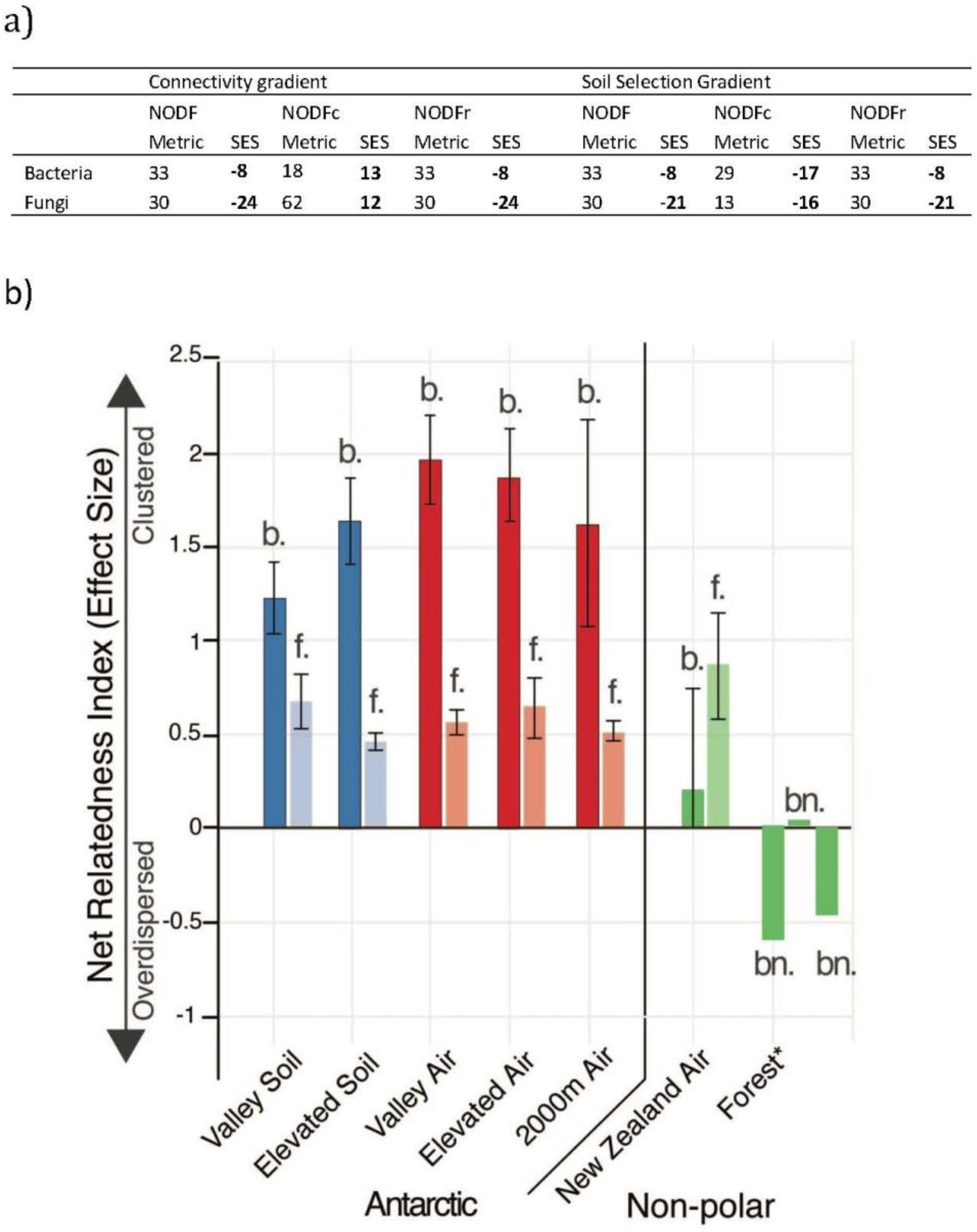
Phylogenetic structuring of local and global pools for bacterial and fungal diversity. **a)** Nestedness estimates made using the NODF model (where 0 = no nestedness, 100 = perfect nestedness). Fungi were more nested (NODFc = 62) than Bacteria (NODFc = 18). Bacteria and Fungi were significantly nested for taxa composition under the hypothesis that nestedness can be maximised by ordering sites from the most connected to the least connected. Least connected sites are demonstrated as a proper subset of richer, more connected sites. **b)** Net Relatedness Index analysis of phylogenetic structure within each sample type for Bacteria (b) and Fungi (f). Error bars show the standard error of the mean for all samples in a given substrate type. Values for highly dispersed non-polar bacterial communities associated with forest soil are given for comparison (bn) and indicated by an asterisk ^62^. Sampling locations: valley (soil and 1.5m above ground), elevated (soil and 1.5m above ground at higher altitude locations at Bull Pass and valley ridges), 2000m (helicopter samples) and New Zealand (non-polar).

The Net Relatedness Index (NRI) added phylogenetic support to the findings of our Network and Nestedness Analysis. NRI analyses demonstrated that local Antarctic communities were not a random sample of the species pool. Antarctic samples of bacteria displayed greater and highly significant phylogenetic clustering than the non-polar samples, which were almost randomly structured and in some case over-dispersed (Fig. 3b). Although the pattern itself does not prove any specific process, the results clearly indicate that Antarctic bacterial communities both in soil and air must have been selected non-randomly, which is consistent with both the taxonomic and phylogenetic observations for our air and soil communities. This result is congruent with observations of biodiversity for other soils in the Dry Valleys region ^31,47–49^. The Fungi were always significantly clustered but bacterial communities were always much more clustered than fungi at any given airborne or soil location. The fungal data should, however, be interpreted with care given current drawbacks with phylogenetic reconstruction based on ITS and despite our efforts to correct for them (Online Supplementary Information: Supplementary Methods). Nonetheless even a cautious interpretation of the data suggests a limited extent of input from fungal taxa not present in local reservoirs. This interpretation concurs with our other lines of evidence presented here and with studies on fungal dispersal from other biomes ^50^.

Contrary to the view that “everything is everywhere” in terms of airborne microbial transport, our data indicates that the aerosphere is a strongly selective habitat that limits dispersal, although the extent may vary between taxonomic groups and spatial scales. We conclude that inter-continental microbial connectivity to the McMurdo Dry Valleys of East Antarctica is limited, and this supports the hypothesis that the Hadley Cell circulation acts as a dispersal barrier to the poles ^17^ even during the austral summer when the Polar Vortex is annually at its weakest ^51^. The Antarctic continent supports other smaller ice-free soil regions and whilst we are unable to directly extrapolate our data to these, it is reasonable to expect similar patterns given what is known of air circulation to the continent; that is, that other Antarctic ice-free areas may also be somewhat decoupled from global microbial reservoirs. An exception may be the peninsula in West Antarctica due to its proximity to the South American continent. Comparison of our soil biodiversity estimates with those for other Dry Valleys locations suggest there is a common core diversity throughout the Antarctic Dry Valleys ^39,47–49,52^. Hence, we expect our data to be broadly applicable to this region. The low level of airborne immigration from exogenous sources may represent an inherently low flux for Antarctica despite supplementation from pulsed inputs of diversity from stochastic events. This may help to explain the unique microbial composition of Antarctic soils compared to others globally ^53^.

We have presented multiple lines of evidence to refute the null hypothesis that local air and soil microorganisms are a random sample of phylogenetic diversity in the regional/global pools. Sources of recruitment other than persistent airborne transport are therefore necessary to fully explain the extant Antarctic soil microbial diversity patterns observed. One potential explanation is stochastic storm events where particulate matter supporting biological propagules is thought to be transported on local scales within the Dry Valleys ^54^, although we did not encounter any such events during our sampling expedition. A further source may be local dispersal from geothermal refugia as they are important reservoirs for radiative dispersal of animal, plant and lichen taxa ^21^. The periodicity from which dispersal from such reservoirs occurs is, however, unknown. An additional reservoir may be the moisture-sufficient soil around lakes where microbial mats are known to persist over inter-annual periods ^55^. Local refugia may be important in facilitating resilience at the landscape scale where severe local extinction pressure occurs due to stochasticity and steep environmental gradients for abiotic variables ^56^. Challenges remain in deciphering the relationship of diversity patterns to biomass and ecosystem function ^32,57^, but the revelation that airborne connectivity is largely localised rather than being an inter-continental scale process emphasises the conservation value of the McMurdo Dry Valleys as a unique ecosystem. This is particularly pertinent in light of a predicted increase in stochasticity for atmospheric air circulation as a result of climate change ^58^, which may led to an increased flux of foreign and invasive taxa into Antarctic ecosystems. Such an increased flux acting in concert with warmer temperatures could profoundly alter the unique biota inhabiting the Antarctic Dry Valleys, one of the of the last pristine ecosystems on Earth.

## Methods

Air mass at near-surface (1.5m above ground) and underlying ultra-oligotrophic soil was sampled from 11 to 23 January 2017 at eight representative valley and elevated locations throughout the Wright Valley, McMurdo Dry Valleys, Antarctica (77.518633 S, 161.768783 E, Supplementary Information Fig. S1). Air mass above the boundary layer for surface influence was achieved by mounting the apparatus in a helicopter with an external sampling port (flightpath: 2,000m AMSL, 77.440836 S, 162.657553 E to 77.524583 S, 161.690917 E). Air sampling was also conducted at the only significant non-polar terrestrial landmass (New Zealand) on the projected back trajectory for incoming air mass. High volume liquid impinger pumps were used to collect airborne microorganisms directly into *RNAlater* nucleic acid preservation solution. Extensive use of controls and apparatus sterilisation ensured high fidelity of the sampling for the ultra-low biomass habitats. Overall, biotic data was retrieved for 30 massive bulk-phase air samples with a total sampled volume of 2,160,000 L and massive air volumes were collected for each discreet air sample (72,000 L). A detailed sampling rationale and methodology is described in the Supplementary Methods (Online Supplementary Information).

Diversity assessments for bacteria and fungi, the two most abundant microbial groups in the Dry Valleys, were made using Illumina MiSeq sequencing of rRNA loci. A total of 3636 bacterial amplicon sequence variants (ASVs) and 5525 fungal operational taxonomic units (OTUs) were identified (Online Supplementary Information: Supplementary Methods). Bacterial 16S rRNA-defined ASVs were delineated using the DADA2 method for exact sample sequence inference ^59^ and fungal ITS-defined OTUs using 97% sequence similarity clustering approach ^60^ (Online Supplementary Information: Supplementary Methods). We achieved near-asymptote in diversity estimation for all samples (Online Supplementary Information: Supplementary Methods). All sequence data generated by this study has been submitted to the EMBL European Nucleotide Archive (ENL) under BioProject PRJEB27416 with accession numbers ERS3573837 to ERS3573946. All diversity metrics and alpha/beta diversity estimates were made using R ^61^ (all packages employed listed in Online Supplementary Information, Supplementary Methods). We employed Network Analysis, Nestedness Analysis and Net Relatedness Index to test the null hypothesis that local air and soil microorganisms are a random sample of phylogenetic diversity in the regional/global pools (Online Supplementary Information: Supplementary Methods).

## Acknowledgements

Field and logistical support was provided by Antarctica New Zealand and the United States Antarctic Program. The authors thank Craig Cary (University of Waikato) for facilitating access to the McMurdo Dry Valleys. The research was funded by a grant from the New Zealand Ministry of Business, Innovation & Employment (UOWX1401) and the Yale-NUS College Start-Up Fund. F.T.M. is supported by the European Research Council (BIODESERT project, ERC Grant agreement n° 647038).

## Author contributions

S.D.J.A. and S.B.P. conceived the study; S.D.J.A and C.K.L. conducted fieldwork; T.M. developed and validated the helicopter sampling method; S.D.J.A. performed laboratory experiments; S.D.J.A., K.C.L., T.C., and S.B.P. performed data analysis and interpretation; D.A.C., F.T.M. and S.B.P. critically assessed and interpreted the findings; S.B.P. wrote the manuscript with input from all authors.

## Materials & Correspondence

Correspondence and requests for materials should be addressed to S.B.P.

## Online Supplementary Information

### Supplementary Methods

To overcome the technological and methodological limitations of earlier approaches to aerosol sampling using low volume air pumps and impactor collection, which are known to introduce significant sampling bias to airborne microbial sampling ^1,2^, we employed a high-volume liquid impinger apparatus (Coriolis μ, Bertin Technologies, France) and developed a novel collection protocol optimised for low-temperature environments. This involved collection of samples directly into RNA*later* nucleic acid preservative solution (Invitrogen, Carlsbad, CA). Evaporation was compensated for using a peristaltic pump set to between 0.5 and 1.3mL per minute with a mixture of phosphate-buffered saline (PBS) and 20% v/v RNA*later* preservative solution. A random collection cone was assembled into the machine but not activated, and these were used as the negative controls. All collection cones were soaked in 1.5% sodium hypochlorite (NaClO) then washed with 70% ethanol and three washes of Milli-Q H_2_0 before being filled with filtered RNA*later*. All sampling equipment was disassembled between locations and cleaned with NaClO, ethanol and Milli-Q H_2_0. Samples in RNA*later* were stored at 4°C during transit from Antarctica and until processed.

Air mass at near-surface (1.5m above ground) and corresponding soil sampling (top 10mm surface soil after removing pebbles and rocks) was conducted from 11 to 23 January 2017 from eight locations throughout the Wright Valley floor (77.518633 S, 161.768783 E) and high elevation locations at the valley ridge and Bull Pass (7.47085 S, 161.77345 E). Air mass above the boundary layer for surface influence was also recovered by mounting the apparatus in a helicopter with an external sampling port (flightpath: 2,000m A>M.S.L., 77.440836 S, 162.657553 E to 77.524583 S, 161.690917 E). Overall, we retrieved 33 massive bulk air samples plus underlying soil samples from 18 locations (high altitude Antarctic air samples and non-polar air did not have accompanying soil samples). Massive bulk-phase air volumes (72,000L per sample) were collected for each discreet air sample location. Biotic data were retrieved for 30 air samples with a total sampled volume of 2,160,000L, and from all soil samples. representing several orders of magnitude increase in sample volume over earlier studies (Kobayashi et al, sample n = 2, approx. 6,000 L total volume, total OTUs identified 213 ^3^; Pearce et al, sample n = 2, total sample volume approx. 370,000 L, total OTUs identified approx. 30 ^4^; Bottos et al, sample n = 2, total volume approx. 150,000 L, total OTUs identified = 202 ^5^).

At each station and time interval air flow rates of 300L/min were employed for 4hrs into 15ml RNA*later*. This approach overcomes limitations from earlier studies where low volume pumps have necessitated long sampling durations with uncertain microbial survival and recovery and impaction techniques that are known to bias against certain phyla ^2,6^. Non-polar air samples were collected from New Zealand’s North Island (36.916153 S, 174.645760 E) during the same austral summer season and using the same method. We selected this place because our HYSPLIT back trajectory analysis (see below) indicated that this was the nearest non-polar land mass from which air mass arriving at the Dry Valleys location was derived. These samples were used to make broad diversity comparisons with possible exogenous sources although we acknowledge that additional variability is likely among non-polar aerosols given inherent uncertainties over their trajectory to the Antarctic. It should be noted that unlike other substrates with high micro-habitat variability such as soil it is neither possible nor necessary to obtain massive numbers of discreet replicate samples for a bulk phase such as air and so our approach focused on obtaining massive volume samples within sampling timeframes of hours which allowed meaningful inference from biological data (rather than pumps operating continuously for weeks or months as in previous studies and with the inherent problems this brings to downstream diversity estimation ^2–5^). Each location was sampled at three discreet time intervals during the austral summer field season and the extremely low variability for diversity estimates from a given substrate support the validity of our approach.

Sampling was conducted within local weather parameters as follows: Relative humidity 20-58%, Temperature −3.3-6.9°C, wind speed 0.6-9.5m/S, wind direction E-ESE (Kestrel 3500 Weather Meter, Nielson-Kellerman Co, Minnesota, USA); Total near-ground air particulate matter 2939-6558μg/m^3^ PM2.5-10 (Aerotrak, TSI Incorporated, Minnesota, USA). Back-trajectories of air mass arriving at each sampling interval were generated using the National Oceanic and Atmospheric Administration (NOAA) HYSPLIT-WEB model (https://ready.arl.noaa.gov/HYSPLIT.php). Separate calculations were made at 3d and 15d because they represent the average minimum and maximum residence times for microorganisms in air transported to the McMurdo Dry Valleys ^7^. HYSPLIT back trajectories were calculated using the GDAS database and the model vertical velocity option. Three day back trajectories travelled between 598 and 2581 km with an average of 1350 km at an average altitude of 2769 m and a maximum altitude of 5174 m above mean sea level (AMSL) Fifteen day back trajectories travelled between 4673 and 11216 km with an average of 6886 km at an average altitude of 3034 m and a maximum altitude of 8211 m AMSL.

Airborne microbial samples from the RNA*later* preservation solution were filtered onto a 25mm 0.2μm polycarbonate filter and stored frozen until processed. Total DNA was directly extracted using a CTAB protocol ^8^. DNA was extracted from three 0.75g ±0.025 of soil using the same CTAB protocol. DNA yield for these ultra-low biomass samples was quantified using the Qubit 2.0 Fluorometer (Invitrogen) in the range 1.06-8.44ng. Samples were then stored at −20°C until processed. We used DNA yield as an indirect estimate for biomass. We were unable to successfully apply direct cell/fluorescent particle counting due to the extremely low cell numbers in Antarctic air, although we acknowledge the validity of this approach in studies of higher biomass aerial habitats ^9^. We did not apply real-time quantitative PCR as a further indirect estimate of biomass since the approach has significant limitations that preclude meaningful estimates of biomass from environmental samples with domain-specific PCR primers ^10,11^.

Illumina MiSeq libraries were prepared as per manufacturer’s protocol (Metagenomic Sequencing Library Preparation Part # 15044223 Rev. B; Illumina, San Diego, CA, USA) and as previously described with PhiX positive controls ^12^. We targeted Bacteria and Fungi since these domains are the most abundant microorganisms in the McMurdo Dry Valleys ^13^. PCR was conducted with primer sets targeting the V3-V4 regions of bacterial and archaeal 16S rRNA gene: PCR1 forward (5’ TCGTCGGCAG CGTCAGATGT GTATAAGAGA CAGCCTACGG GNGGCWGCAG 3’) and PCR1 reverse (5’ GTCTCGTGGG CTCGGAGATG TGTATAAGAG ACAGGACTAC HVGGGTATCT AATCC 3’) and the internal transcribed spacer region of fungal 18S and 5.8S rRNA genes: ITS1 forward (5’ CTTGGTCATTTAGAGGAAGTAA 3’) and ITS2 reverse (5’ GCTGCGTTCTTCATCGATGC 3’). These primers for Bacteria and Fungi are widely accepted to capture the broadest estimates of diversity ^14–16^ and were used according to recommended workflows for the Earth Microbiome Project (http://www.earthmicrobiome.org). Total sequence library sizes were 3,994,561 for bacteria and 2,437,256 for fungi before filtering and total counts in the processed dataset were 1,333,553 bacterial and 2,220,883 fungal sequences. A total of 3,636 bacterial and 5,525 fungal taxa were identified from these. All sequence data generated by this study has been submitted to the NCBI Sequence Read Archive under BioProject PRJEB27416 with accession numbers ERS3573837 to ERS3573946.

Sequencing data for 16S rRNA gene amplicons was processed based on the DADA2 v1.8 ^17^ pipeline. Primers sequences were removed using cutadapt ^18^ to remove forward (CCTACGGGNGGCWGCAG) and reverse (GACTACHVGGGTATCTAATCC). The reads were uniformly trimmed to 280 bp (forward) and 250 bp (reverse) and then filtered by removing reads exceeding maximum expected error of 2 for forward reads and 5 for reverse reads or reads containing ambiguity N symbol. The reads were used to train the error model and then dereplicated to acquire unique sequences, which were used to infer sequence variants with the trained error model. The forward and reverse reads were merged and chimeric sequences were removed. For bacteria we used amplicon sequence variants (ASV) to assign operational taxonomic units (OTUs) since this has been shown as the most robust method currently available for bacterial 16S rRNA gene-defined taxa identification ^19^. OTUs were given taxonomic assignment using DADA2 with SILVA nr v132 database ^20^ to provide species level assignment based on exact match between ASVs and known reference sequences. For fungal ITS1 amplicon data, the sequences were processed using USEARCH v9.0.2132 ^21^. The forward and reverse paired-end sequences were merged and filtered to remove reads >1 maximum expected error per sequence. Additionally, anomalous sequences (<200 or >500 bp in length or exceeding 20 homopolymers) were also removed. After dereplication and removal of singletons, the reads were clustered at 97% identity threshold to obtain representative sequences as OTUs ^14^. Unfiltered reads were mapped onto these OTUs to produce an abundance table of the occurrence of these OTUs within the communities. The representative sequences were given taxonomic assignment using USEARCH SINTAX classifier and RDP Warcup training set v2 (rdp_its_v2) ^22^.

The resulting OTUs were then processed as previously described ^23^. The R packages phyloseq ^24^, DESeq2 ^25^ and ggplot2 ^26^ were used for downstream analysis and visualisation including ordination and alpha/beta diversity calculations. Despite inherent bias due to underlying differences in substrate biomass influenicng species richness estimates with any cross-habitat biogeographic analysis ^27^, we are confident that comparable yet inherently low biomass in all our air and ultra-oligotrophic mineral soil samples minimised such influence and this was reflected in our diversity estimates. An exception was that fungi in non-polar air were markedly more taxon-rich than in Antarctic samples. Therefore we used guild analysis (FUNGuild ^28^) to establish the predominantly phyllosphere origin of fungi in non-polar samples which are absent in Antarctica as well as the latitudinal gradient in fungal diversity ^29^ (lack of database depth for FUNGuild limited its value to identifying ecological guilds for non-polar fungi only). For heatmap visualisations the 1,000 most abundant OTUs in each data set were selected, and these captured 91% bacterial and 96% fungal sequences. All other analysis used the entire sequence library data. Our heatmap analysis therefore had high confidence since the unsampled ‘tail’ comprised only extremely rare sequence variants at very low/singleton abundance.

We used multiple statistical approaches to test the null hypothesis that local soil and air sample communities were a random sample of the of the regional pool. Rejection of the hypothesis (i.e., non-random patterns) yields observational evidence for the alternative hypothesis that local communities are a non-random selection from the regional pool. Specifically, we expected local soil communities to be selected against the extreme conditions found at soils in the McMurdo Dry Valleys soil and thus to display clustering. At the same time, we expected air samples to be less structured or even random and thus better reflect the regional pool, although some structuring due to local influences from atmospheric stressors such as low temperatures and UV exposure were also be expected.

We employed approaches that utilised both taxonomic identity and phylogenetic structure of the communities. Ecological Network Analysis is a commonly employed tool to infer biotic interactions within and between communities by visualising links between species nodes. Potential relationships pertinent to our system include connectivity, clustering and nestedness which are informative to interpreting the biogeographic patterns of species occurrence ^30^. We performed Ecological Network Analysis on our samples using the R package phyloseq ^24^ with maximum Bray-Curtis distance of 0.2 for Bacteria and 0.4 for Fungi to establish connection between nodes (representing communities). The nodes were positioned using the Fruchterman-Reingold method ^31^.

Nestedness is a widespread biogeographical pattern that emerges when species composition of small assemblages is a nested subset of larger regional assemblages. We quantified this pattern using the metrics NODF ^32^ and its compositional (NODFc) and incidence (NODFr) version. This metric is currently considered one of the most effective and statistically robust, especially in relation to the null models that are used to test whether observed metrics are smaller or larger than expected by chance ^30,33^. The metric ranges from 0, that is no nestedness or perfect antinestedness, to 100 (perfect nestedness). In a perfectly nested assemblage, species poor communities are just a subset of species richer communities, or less frequent species always occur in subsets of sites where most widespread species also occur, or a combination of both. We applied the NODF metrics to the matrix of the investigated sites both for Bacteria and Fungi. We simulated null models using a random swap algorithm (R project, vegan package, function “oecosimu” ^34^) with fixed row and column sums. This combination to create a random matrix is the most conservative in terms of Type I and II errors and is particularly recommended when species co-occurrence is critical to the tested hypothesis ^30^. We calculated Standardised Effect Sizes and tested for significance of effects with 999 permutations and at P < 0.01. We calculated NODF and null models by a specific order of sites in the matrix that reflected our main null hypothesis. This was that local communities are just a passive sampling from regional species pools. Specifically, we used two complementary orderings of the sites to test our hypothesis. First, we created a connectivity gradient, which assumed New Zealand is the site more connected to the global species pool while soils in the Antarctic Dry Valleys are the least connected (the exact order was: non-polar>high-altitude air>elevated air>valley air>elevated soil>valley soil). In the second test of the hypothesis local, more isolated soils were assumed to select more than connected soil and air habitats. In this case, sites were ordered as follows: Valley Soil>Elevated Soil>Valley Air>Elevated Air>high-altitude air>non-polar air.

Community phylogenetic metrics were calculated using the R ^35^ packages picante ^36^, ape ^37^, phylobase ^38^, adephylo ^39^, and phytools ^40^. Phylogenetic trees for community phylogenetic structure analysis were constructed for all OTUs with FastTree v2.1.9 on multiple alignment of sequences produced by MUSCLE v3.8.31. For Bacteria an approximately Maximum-Likelihood approach was used whilst for Fungi an alignment-free distance approach with Neighbour-Joining method was employed in order generate a ITS-based phylogenetic tree for community metrics ^41^. The distance approach was used for Fungi as the hypervariability of ITS1 loci hinders multiple sequence alignment required for most phylogenetic analyses. To validate this approach, we compared our tree topology with the most recent whole genome phylogenies for the Fungi ^42,43^. This approach was robust at higher taxonomic levels (there is no consensus for fungal phylogenies using multiple loci or whole genomes below Order rank) and has been used successfully with other eukaryotic taxa as a workflow for Net Relatedness analysis ^44^. Although we acknowledge limitations to this approach, the advantages of using ITS loci for taxonomic identification vastly outweighed its shortcomings, and we also triangulated data from this test with additional analytical approaches, so overall our inclusion of this test is justified and interpreted conservatively. Mean phylogenetic distance (MPD) ^45^ was calculated to measure phylogenetic distance between ASVs and OTUs in each sample and converted to the Net Relatedness Index (NRI) by multiplying them by −1. A null model algorithm based on independent swap (999 randomisation) was used to test the extent of phylogenetically clustering (positive values) or overdispersion (negative values) ^46^. Results for NRI were expressed as effects size, (MPD –MPDnull)/SD(MPDnull)]. Distance Decay of phylogenetic versus geographic distance for bacteria and fungi was estimated using the R package Vegan ^34^.

---

**Fig. S1.**
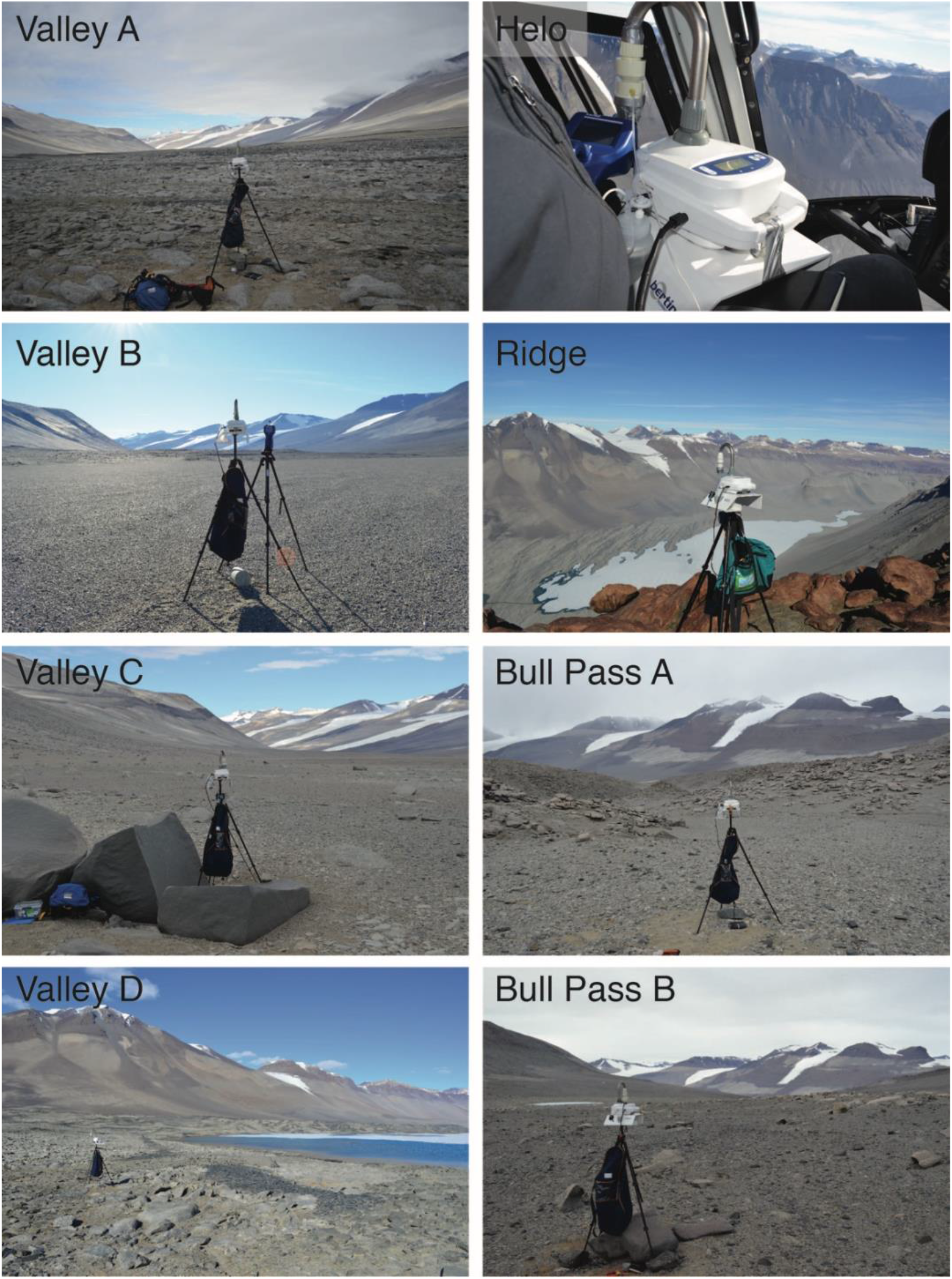
Sampling locations in the Wright Valley, McMurdo Dry Valleys, Antarctica. All ground sampling was completed at a consistent height at a site removed from significant topographical variation to ensure minimal local wind effects. The air intake was orientated towards the prevailing wind direction at the time of sampling and was only approached from downwind. Helicopter sampling (Helo) was conducted at 2,000 m AMSL with the intake outside of the passenger window.

**Supplementary Figure S2.**
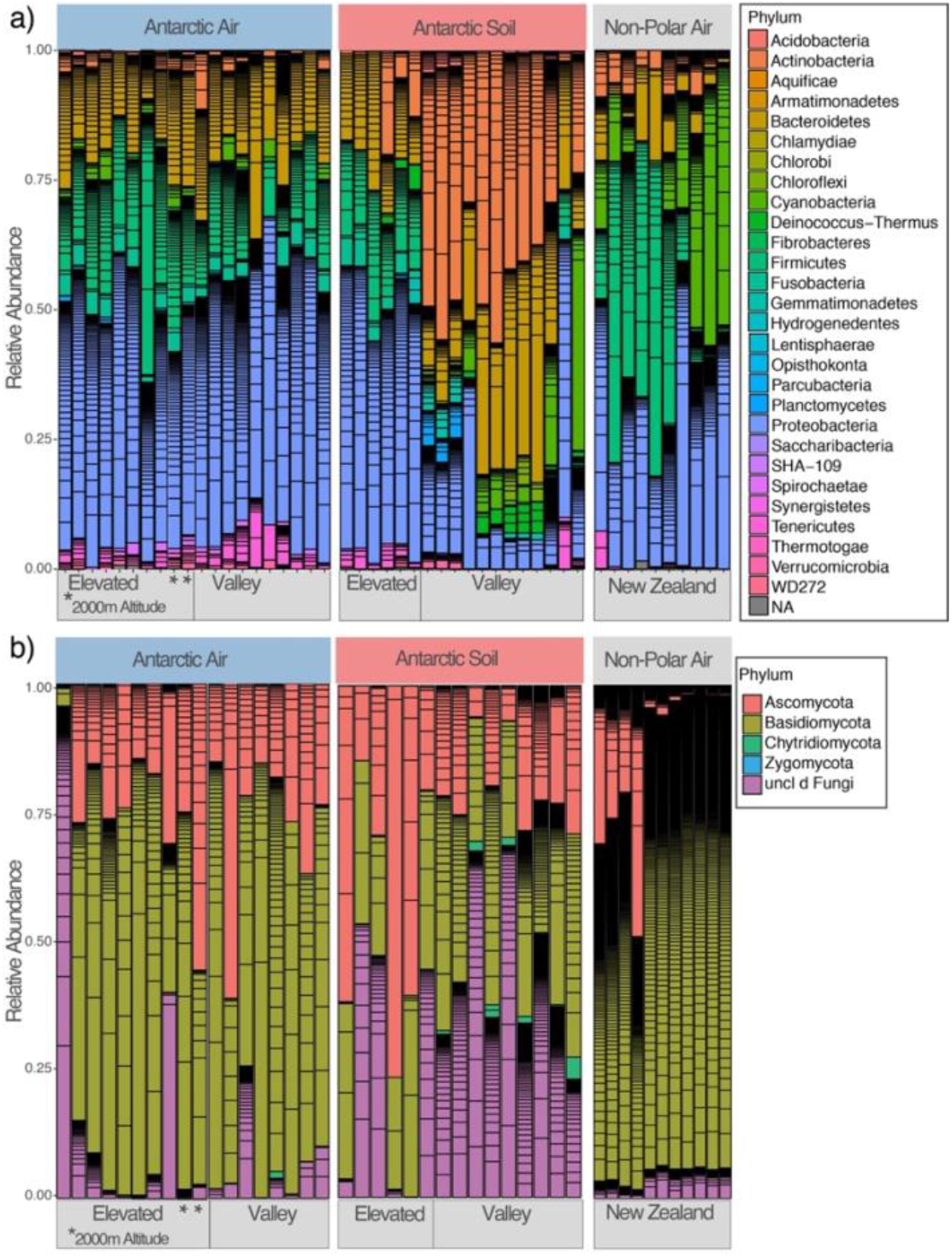
Interactive bar charts for taxonomic identity of bacteria and fungi in each air and soil sample. Individual charts are presented for taxonomic identity at Phylum, Class and Genus. Readers can select any item in the plots and the taxon identity and relative abundance will be displayed. The overall bar charts for a) Bacteria and b) Fungi for all Antarctic and non-polar samples are displayed here at the Phylum level for reference. Black lines indicate delineations at lower taxonomic ranks. Bacteria in non-polar air displayed similar taxa richness to Antarctic air whereas Fungi were markedly more diverse and Guild Analysis was employed to illustrate this reflected phyllosphere input ^28^. File names for interactive bar charts:

1. Bacteria_Phylum.html
2. Bacteria_Class.html
3. Bacteria_Genus.html
4. Fungi_Phylum.html
5. Fungi_Class.html
6. Fungi_Genus.html

